# Slow Intrinsic Oscillations in the Ventrolateral Preoptic nucleus

**DOI:** 10.1101/2025.10.13.682168

**Authors:** Quentin Perrenoud, Jérôme Ribot, Hélène Geoffroy, Thierry Gallopin, Nathalie Rouach, Armelle Rancillac

**Affiliations:** Department of Neuroscience, Yale School of Medicine, Kavli Institute for Neuroscience, Wu Tsai Institute, New Haven, CT, United States; Neuroglial Interactions in Cerebral Physiology and Pathologies, CIRB, Collège de France, CNRS UMR 7241, INSERM U1050, PSL University, PSL-Neuro, Paris, 75005, France; Brain Plasticity Unit, ESPCI-Paris, PSL Research University, Paris, F-75005, France

**Keywords:** Hypothalamus, Sleep-promoting neuron, noradrenaline, intrinsic bursting, UP and DOWN states, oscillation, rhythm

## Abstract

The ventrolateral preoptic nucleus (VLPO) of the hypothalamus plays a major role in the induction and consolidation of non-rapid eye movement (NREM) sleep. While VLPO neurons are heterogeneous, they often display low-threshold spikes (LTS), a feature that supports rhythmic activity. Nevertheless, rhythmic bursting in these VLPO neurons has never been observed. Here, we report that ∼12% of VLPO neurons in a large database of *ex vivo* patch-clamp recordings display spontaneous rhythmic bursting of action potentials. This activity occurred in putative sleep-promoting neurons, identified by inhibitory responses to noradrenaline (NA), as well as wake-active neurons that were activated by NA. Unsupervised clustering of 24 neurons based on burst properties, electrophysiological, and morphological features revealed three distinct groups: one corresponding to putative sleep-promoting neurons and two wake-active neurons with fast and slow bursting dynamics. Strikingly, membrane potential oscillations persisted in the presence of tetrodotoxin (TTX), indicating that rhythmic bursting is driven by intrinsic mechanisms rather than network activity. These findings suggest that rhythmic bursting is an intrinsic and functionally relevant mode of activity in VLPO neurons, which may contribute to sleep regulation.

## Introduction

Sleep is a fundamental and ubiquitous behavior across the animal kingdom, but its function and mechanism remain actively debated (Shein-Idelson et al., 2016). In mammals, sleep alternates between two distinct physiological states: Non-Rapid Eye Movement (NREM) sleep and Rapid Eye Movement (REM) sleep. The induction and maintenance are regulated by complex networks of interconnected brain nuclei spanning the forebrain, midbrain and hindbrain (Brown et al., 2012; Kashiwagi et al., 2024; Scammell et al., 2017; Weber and Dan, 2016). Among these, the ventrolateral preoptic nucleus (VLPO), a diffuse region of the ventral hypothalamus, constitutes a key promoter of NREM sleep (Arrigoni and Fuller, 2022; Reitz and Kelz, 2021). However, the circuit mechanisms underlying its function are not yet fully understood.

The VLPO receives convergent inputs from several wake-promoting regions, including the locus coeruleus (LC), the tuberomammillary nucleus (TMN), and the lateral hypothalamus (LH), which respectively release noradrenaline (NA), histamine, and orexin. It also integrates cholinergic, serotonergic, glutamatergic, and GABAergic projections arising from various brain sites (Chou et al., 2002). Among the heterogeneous neuronal populations in the VLPO (Dubourget et al., 2017; Gallopin et al., 2005, 2000; Sangare et al., 2016), a key subset, referred to as sleep-promoting neurons, is directly inhibited by NA (Gallopin et al., 2000; Liang et al., 2021) and indirectly inhibited by histamine (Liu et al., 2010; Sherin et al., 1998; Williams et al., 2014) and orexin (De Luca et al., 2022, 2017). These neurons, in turn, inhibit wake-promoting nuclei via GABAergic and galaninergic projections (Kroeger et al., 2018; Sherin et al., 1998, 1996), thereby contributing to sleep onset and maintenance.

A commonly proposed model suggests that the mutual inhibition between wake-promoting centers and sleep-promoting VLPO neurons creates a bistable system that toggles between sleep and wakefulness (Saper et al., 2010). Nevertheless, this model does not fully account for several observations. Notably, simple mutually inhibitory circuits typically generate abrupt and stochastic state transitions, whereas natural transitions between wakefulness, NREM and REM sleep are gradual and well ordered, suggesting the presence of complex regulatory mechanisms (Alam et al., 1999; Chou et al., 2002; Gallopin et al., 2005; Kroeger et al., 2018; Scharbarg et al., 2016; Christophe Varin et al., 2015). This highlights a need for further research aiming at understanding the mechanisms governing the patterning of activity in VLPO neurons.

Neural types other than sleep-promoting neurons exist in the VLPO (Dubourget et al., 2017; Gallopin et al., 2005, 2000; Sangare et al., 2016). These neurons are predominantly GABAergic and excited by NA, suggesting they may exert inhibitory control over sleep-promoting neurons during wakefulness (De Luca et al., 2017; Liu et al., 2010; Williams et al., 2014). However, their precise contribution to sleep-wake regulation remains unclear. In addition, VLPO neurons are often capable of emitting low-threshold spike (LTS) (Dubourget et al., 2017; Gallopin et al., 2000, 2005; Scharbarg et al., 2016; Christophe Varin et al., 2015), a rare electrophysiological property that enables the maintenance of oscillation through a depolarizing rebound after hyperpolarization (Crunelli et al., 2005, 1989). Yet, whether VLPO neurons exhibit such oscillatory activity, and how it contributes to their role in sleep regulation, remains unknown.

Here, we report that a subset of VLPO neurons exhibits intrinsically-generated, slow rhythmic bursting activity. Among a dataset of 393 *ex vivo* recordings (cell-attached and whole-cell), we identified 37 neurons displaying spontaneous rhythmic bursting. Twelve of them were inhibited by NA, a hallmark of sleep-promoting identity. We further characterized the electrophysiological and morphological properties of 24 bursting neurons recorded in the whole-cell configuration. Using unsupervised clustering, we identified three different neuronal subtypes: two groups of NA-activated neurons exhibiting rapid and slow bursting activity, respectively, and a third group of NA-inhibited (putative sleep-promoting) neurons with intermediate bursting frequencies. In a subset of 8 neurons, the application of tetrodotoxin (TTX) abolished bursting in only two neurons, whereas it did not affect the oscillations of the membrane potential in the remaining 6 neurons. Together, these findings reveal that VLPO neurons can intrinsically generate rhythmic activity across subtypes, which might contribute to NREM sleep-related rhythmicity.

## Materials and Methods

### Animals

Male-only C57BL/6J mice (14–21 days old; Charles River, France) were housed in a temperature-controlled (20–22°C) room under a 12–hour light-dark cycle (lights on at 09:00 a.m.) with *ad libitum* access to food and water and acclimated in the laboratory for at least 1 week before experiments. All animal procedures were conducted in strict compliance with institutional protocols and were approved by the European Community Council Directive of 22 September 2010 (010/63/UE) and the local ethics committee (Comité d’éthique en matière d’expérimentation animale number 59, C2EA—59, ‘Paris Centre et Sud’). The number of animals in our study was accordingly kept to the necessary minimum.

### Preparation of acute hypothalamic slices

Animals were decapitated at the beginning of the light phase, between 09:00 and 10:00 a.m. Brains were quickly extracted and submerged in ice-cold artificial cerebrospinal fluid (aCSF (mM): 130 NaCl; 5 KCl; 2.4 CaCl_2_; 20 NaHCO_3_; 1.25 KH_2_PO_4_; 1.3 MgSO_4_; 10 D-glucose; 15 sucrose, pH = 7.35) containing the glutamate receptor blocker kynurenic acid (1 mM) and constantly oxygenated with 95% O_2_–5% CO_2_. Coronal brain slices (300 µm thick) were cut with a vibratome (VT2000S; Leica) and stored in oxygenated aCSF containing kynurenic acid until further use.

### Loose patch and whole-cell patch-clamp recordings

Individual slices were placed on the stage of an upright microscope (Zeiss) in a recording chamber maintained at 32°C and continuously superfused with oxygenated artificial cerebrospinal fluid (aCSF). Slices were visualized under infrared light with Dodt gradient contrast (Luigs & Neumann) through a 40X immersion objective and a CCD camera (Hamamatsu). Electrophysiological traces were amplified with a MultiClamp700B amplifier (Axon Instruments) and digitized by an acquisition board (Digidata 1440; Axon Instruments) connected to a computer running pCLAMP (Axon Instruments). Recordings were targeted to the VLPO and performed in the whole-cell and loose cell-attached configurations with patch-clamp pipettes (3–6 MΩ) filled with an internal solution containing (in mM): 144 K-gluconate; 1 MgCl_2_; 0.5 EGTA; 10 HEPES (pH 7.2) and 2 mg/mL biocytin (Sigma Aldrich) for morphological analysis. The osmolarity of the internal solution was adjusted to 285–295 mOsm. The liquid junction potential of the patch pipette and the perfused extracellular solution were estimated at 11 mV and were not corrected.

Spontaneous currents were recorded in voltage-clamp mode at a holding potential of **-**60 mV, which is more depolarized than the reversal potential for inhibitory postsynaptic currents (IPSCs). The pipette solution contained a low chloride concentration (1 mM), resulting in an E_Cl_ of-111 mV. Therefore, inward currents were identified as spontaneous excitatory postsynaptic currents (sEPSCs) and outward currents as sIPSCs. Signals were filtered at 2 kHz, digitized at 10 kHz and acquired online using the pCLAMP 9 (Clampex; Axon Instruments).

### Drugs delivery

Noradrenaline (NA, 10 µM; Sigma), serotonin (5-HT, 100 µM; Sigma) or Tetrodotoxin (TTX, 1 µM, Sigma) were mixed in aCSF and applied to slices via superfusion during 30 s. Drug application was preceded and followed by superfusion with regular aCSF.

### Histology

After recordings, slices were immersed overnight in 0.1% phosphate buffer (PB) adjusted to pH 7.4 containing 4% paraformaldehyde and stored in PB at 4°C until further use. Biocytin was revealed through diaminobenzidine staining (DAB, Vector) performed after binding of avidin-horseradish peroxidase through kit ABC (Thermofisher). Slices were mounted in MOWIOL.

### Burst analysis

Action potentials were detected using the template search function of Clampfit (pCLAMP10, Molecular Probes). Bursts of action potentials were then identified using the Clampfit “Poisson Surprise” protocol (Legendy and Salcman, 1985) and characterized by the following parameters. The average (1) **up-state** and (2) **down-state** membrane potential were measured, and (3) **delta** was taken as their difference. Spike train autocorrelograms were computed for each recorded neuron using a symmetrical time window ranging from –2500 ms to +2500 ms with 20 ms bin resolution. To avoid contamination by the large peak around 0 ms corresponding to the refractory period and immediate spike correlations, the central window from –250 ms to +250 ms was systematically excluded from further analysis. To quantify rhythmicity, a sine wave of the form y(t) = a sin(2πf t) +b, where a is the oscillation amplitude, f the frequency, and b the vertical offset, was fitted to the remaining autocorrelogram data using a least-squares optimization procedure. (4) The rhythmicity index was defined as the **ratio a/b**, where higher values reflect stronger oscillatory modulation relative to baseline autocorrelogram level, and (5) the **frequency f=1/T** corresponds to the dominant rhythmic firing frequency (respectively period T) of the neuron. (6) **Rise time** and (7) **decay time** were measured in Clampfit with cursors manually placed at the beginning and end of state transitions and averaged over bursts. Likewise, (8) **burst duration** and (9) **number of action potentials** per burst were averaged over bursts for each recording. (10) **Intra-burst frequency** and (11) **inter-burst frequency** were the number of action potentials per unit of time during and outside of bursts, respectively. (12) The **overall firing rate** and (13) **mean inter-burst intervals** were also measured.

### Electrophysiological properties

In whole-cell recordings, 27 electrophysiological parameters were measured from the voltage responses induced by 800 ms current steps, ranging from - 100 pA to firing saturation in 10 pA increments, while neurons were at a membrane potential of-60 mV (Ascoli et al., 2008; Karagiannis et al., 2009). (1) The Resting Membrane Potential (**RMP**) was measured immediately after achieving whole-cell configuration. (2) Input resistance (**Rm**) was estimated from the voltage deflection elicited by-10 pA hyperpolarizing current using Ohms law (R = U/I). (3) Membrane time constant (**τm**) was measured on the same trace as the time constant of an exponential fitted to the response onset. (4) The membrane capacitance (**Cm**) was calculated according to the equation Cm = τm/Rm. Under our conditions, injection of hyperpolarizing current pulses often induced a hyperpolarization-activated cationic current (Ih) that followed the initial hyperpolarization peak, known as a sag. (5) **R_hyp_** and (6) **R_sag_** were measured as the slope of the linear portion of an I–V plot measured at the beginning (0−0.1 s;-100 to 0 pA) and at the end of hyperpolarizing current pulses (0.7– 0.8 s;-100 to 0 pA). (7) **ΔG_Sag_** corresponds to (R_sag_**–**R_hyp_)/R_sag_. A distinctive electrophysiological feature of sleep-promoting VLPO neurons is the presence of Low Threshold Spikes (LTS), i.e., a burst of action potentials riding on a calcium depolarizing wave that occurs on rebound from hyperpolarization. (8) **Post-inhibitory rebound** (with or without action potential) following hyperpolarizing current of 100 pA from-60 mV and (9) of **LTS** (rebound and spike) following current pulses resulting in depolarization inferior to-80 mV were encoded as binary variable. (10) **Rheobase** was the minimum current eliciting an action potential. (11) The **first spike latency** was the time needed to elicit a spike at rheobase from current injection onset. Neurons can exhibit bursting, adapting, regular, or stuttering firing behavior in response to depolarizing currents. To capture this diversity, the instantaneous firing frequency (1/inter-spike intervals) was measured in response to the minimal current injection eliciting more than three action potentials and fitted with a linear curve. (12) Adaptation (**m_threshold_**) and (13) minimal steady state frequency (**F_threshold_**) were taken respectively as the slope and the intersect of this linear fit. Instantaneous firing frequency was then measured on the maximal current step before saturation and fitted with the sum of an exponential and a linear function according to the equation F_sat_ = A_sat_ × E^−t/τsat^ +t × m_sat_ + F_max_, where (14) **A_sat_** corresponds to the amplitude of early frequency adaptation, (15) **τ_sat_** to the time constant of early adaptation, (16) **F_max_** the maximal steady-state frequency, and (17) **m_sat_** to the slope of late adaptation. (18 and 24). Amplitudes of first (**A1**) and second (**A2**) action potentials were measured from spike onset in the train induced by the minimal current step evoking at least 3 action potentials, and (19 and 25) spike durations (**D1** and **D2**) were measured on the same spikes at half-amplitude. (20) The amplitude (**AHP_max_**) and (21) time (**tAHP_max_**) of after-hyperpolarization were measured relative to spike onset on the first action potential. When AHPs presented a biphasic shape, (22) the amplitude (**ADP**) and (23) time (**tADP**) of the rebound (i.e., the so-called After Depolarization Potential) were measured from the first hyperpolarization peak. (26) Amplitude variation (**Var A**) and (27) duration variation (**Var D**) were calculated as (A1-A2)/A1*100 and (D2-D1)/D1*100, respectively. All neurons responded to NA. In addition to intrinsic electrophysiological properties (28), the hyperpolarizing or depolarizing pharmacological response to NA application was recorded in current-clamp mode and encoded as either 0 or 1, respectively, by a binary variable.

### Morphological analysis

Neuron morphology was quantified with 10 parameters related to the features of somata and 24 parameters related to those of dendrites. Morphologies of somata were reconstructed from infrared images taken before whole-cell recording using Image-Pro 7 software (Media Cybernetics Inc.). Morphological features were measured from these images after calibration with a standard 24 µm grid. The soma contour was manually drawn from the image, and (1) the cell body **area** and (2) **perimeter** were measured. (3) The **form factor** was defined as the ratio between the perimeter and the area. The (4) **maximal diameter** and (5) **minimal diameter** passing through the centroid were measured, and (6) the **aspect ratio** was taken as the maximal over the minimal diameter. (7) The **somatic solidity** was defined as the ratio of the somata area over its convex area. Likewise, (8) **convexity** was taken as the ratio of the convex perimeter over the perimeter. The measurement of how closely this shape approached that of a circle was assessed throughout (9) the roundness. Finally, (10) somatic compactness was defined as √(4/π) × area/D_feret_max_ where D_feret_max_ is the maximal feret diameter (i.e., the diameter as if measured by calipers).

The dendritic properties were measured from 3-dimensional reconstructions of biocytin-filled arborization performed using the Neurolucida platform. Parameters were extracted using Neurolucida Explorer (MBF Bioscience) and Excel (Microsoft). To be able to describe the differences of dendritic arborizations, we systematically quantified the following parameters:

(1) Number of primary dendrites, (2) total dendritic length, (3) total dendritic surface, (4) total dendritic volume, (5) ratio of the dendritic length over dendritic surface, (6) highest dendritic order segment, (7) average tortuosity of dendritic segments (defined as the ratio of the length of the segment over the straight line path between its extremities), (8) number of dendritic nodes and (9) dendritic planar angle. We performed a Sholl analysis (Sholl, 1953) to describe differences between neuronal arborization. (10) The length of the dendritic arborization that could be enclosed within a 100 μm radius circle around the cell body, (11) between 100 μm and 200 μm radii, (12) between 200 μm and 300 μm radii, and (13) outside a 300 μm radius was systematically extracted. (14) Kdim, a measure from fractal analyses, represented the degree to which the dendritic arbor has a scale-invariant topology. Finally, (15-23) the number of dendritic spines in segment order, ranging from 1 to 8, was extracted, and (24) mean spine density was calculated.

### Ward’s Clustering and Silhouette

Unsupervised clustering of bursting neurons was performed in the matlab environment (Mathwork) as described previously (Dubourget et al., 2017; Hardy et al., 2021; Karagiannis et al., 2009; Q Perrenoud et al., 2012) and was based on a total of 34 parameters: **nine burst parameters**: Up State, Down State,, Delta, a/b, Rise Time, Decay Time, Burst duration, Burst Frequency and Interevent interval; **2 somatic morphological parameters**: cell body area and perimeter and **23 electrophysiological parameters**: RM, Rm, tIC, Cm, Rhyp, Rsag, ΔG_Sag_, LTS, Rebond, Rheobase, 1st spike delay, Asat, tsat, Csat, A1, D1, AHP max, ADP1, tAHP, tADP1, Var A, Var D, NA.

Briefly, parameters were z-scored and a dendrogram of neuron’s proximity in the parameter space was constructed with Ward’s method (Ward, 1963) based on Euclidean distances. Clusters generated by Ward’s method were further refined using the K-means algorithm initialized on Ward’s cluster centroid to ensure that clusters did not overlap. The quality of a clustering was assessed to determine the optimal clustering number (Shin et al. 2013) using silhouette analysis (Rousseeuw, 1987) and the sum of squared errors (SSE).

Silhouette value was defined for each neuron *i* as:

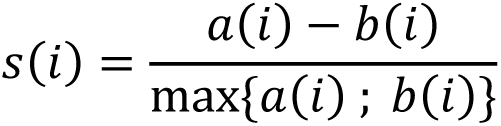

Where *a(i)* is the average distance of neuron *i* to the neurons from its cluster, and *b(i)* is the average distance to the neurons of the closest cluster. A positive silhouette value indicates that, on average, the neuron is closer to the neurons of its own cluster than to the neurons belonging to the closest clusters. On the contrary, a negative value indicates a potential misclassification.

SSE was defined as

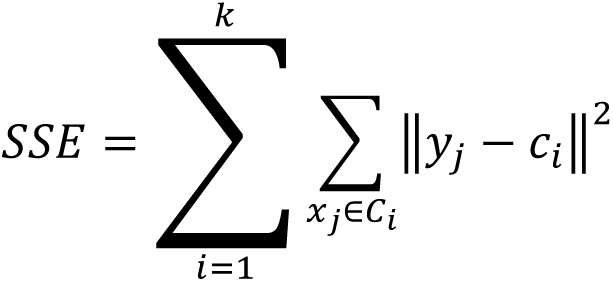

Where *y_j_* is the *j*th object in cluster *C_i_* and *c_i_* the center of cluster *C_i_*.

*Statistics*. All data are expressed as the mean ± standard error of the mean (SEM). Statistical differences between the three clusters were determined using the Kruskal-Wallis one-way nonparametric ANOVA. *P*-values of ≤ 0.05 were considered statistically significant. In all cases, n refers to the number of examined neurons.

## Results

We investigate the properties of VLPO neurons with patch-clamp recording in juvenile slices, and occasionally observed spontaneous, rhythmic activity in the form of a regular burst of action potentials (AP). We thus explored the properties of bursting VLPO neurons.

From a random sampling of VLPO neurons in our previously published patch-clamp datasets (Dubourget et al., 2017; Sangare et al., 2016; Scharbarg et al., 2016; Varin et al., 2015), we analyzed 393 recordings and identified 49 spontaneously bursting neurons, accounting for 12.5% of the total (Fig. 1A). VLPO neurons typically respond to bath application of NA either with inhibitory hyperpolarization (NA(-) neurons) or excitatory depolarization (NA(+) neurons), NA(-) neurons corresponding to putative sleep-promoting neurons (Gallopin et al., 2000). Given the relevance of NA responsiveness to VLPO physiology, we restricted analyses to recordings in which NA responses were assessed (Materials and Methods), yielding a final dataset of 37 VLPO neurons recorded in the loose patch (n = 18) and/or whole-cell configuration (n = 24). For five neurons, recordings were obtained sequentially in both configurations, and in all instances, the shift to a whole-cell configuration did not affect bursting.

**Figure 1.**
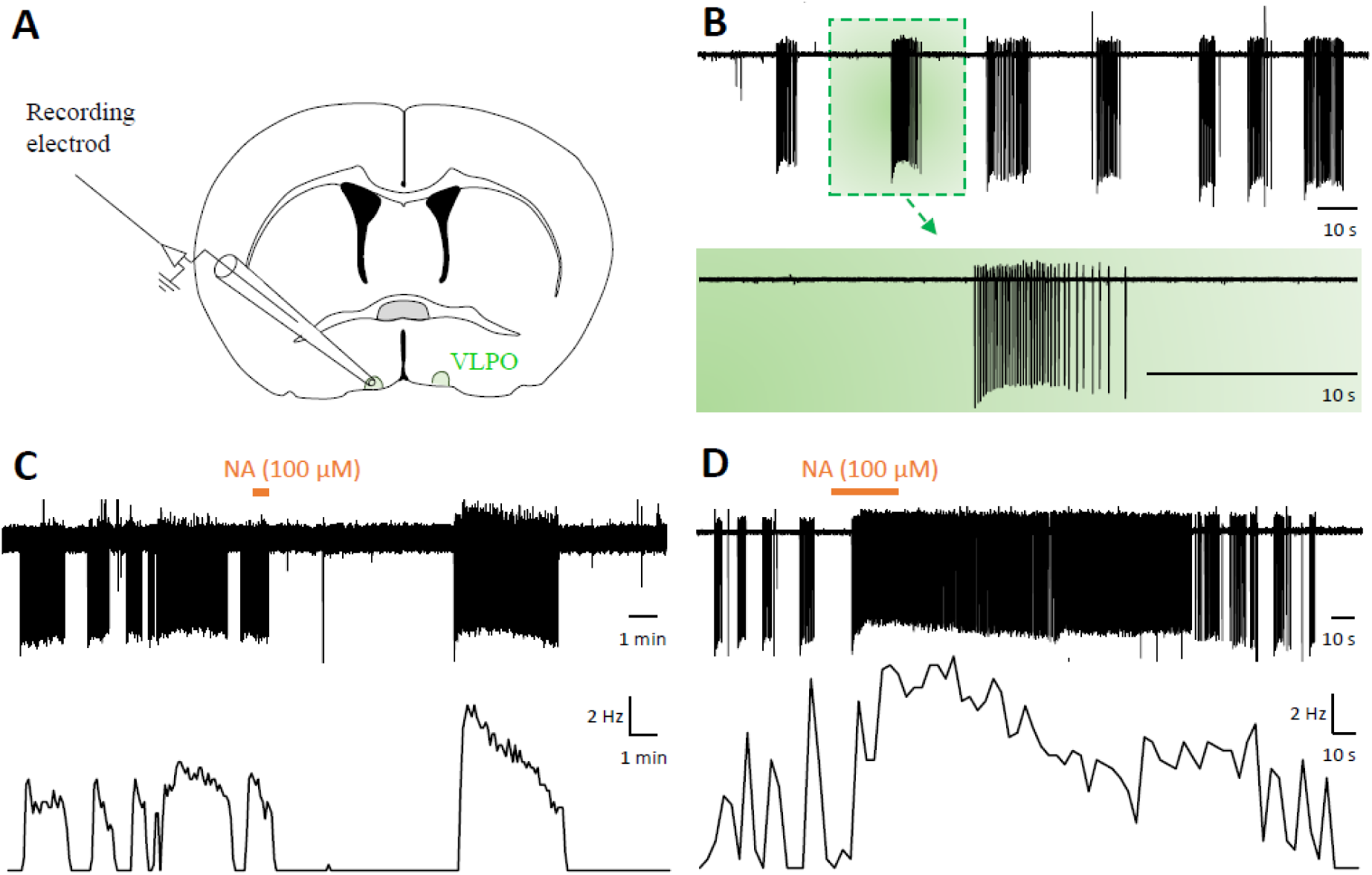
Recordings of bursting neurons in loose-patch configuration in the VLPO. A, Localization of the VLPO and the recording electrode on a coronal slice (around bregma 0.18). **B**, Typical recording of rhythmic sequences of discharge in loose cell-attach configuration within the VLPO (upper trace). An expansion of one bursting sequence is represented in the lower panel. **C** and **D**, NA (10 µM) robustly decreases (C) or increases (D) the firing frequency of discharge. Discharge rates are represented in the lower trace with a bin of 5 s.

We first investigated the properties of neurons recorded in the loose patch configuration. Neurons from this sample were either inhibited (n = 7/18; Fig. 1C) or excited (n = 11/18; Fig. 1D) by NA application. A detailed comparison of the bursting features of NA(-) and NA(+) neurons (Table 1) revealed that NA(-) exhibited a longer burst duration and a lower burst frequency compared to NA(+) neurons. However, the mean action potential (AP) number per burst, intra-burst frequency, inter-burst frequency, and mean AP frequency showed no differences. Overall, these results indicated that bursting in the VLPO is expressed across neuronal subtypes albeit with seemingly variable dynamics.

**Table 1.**
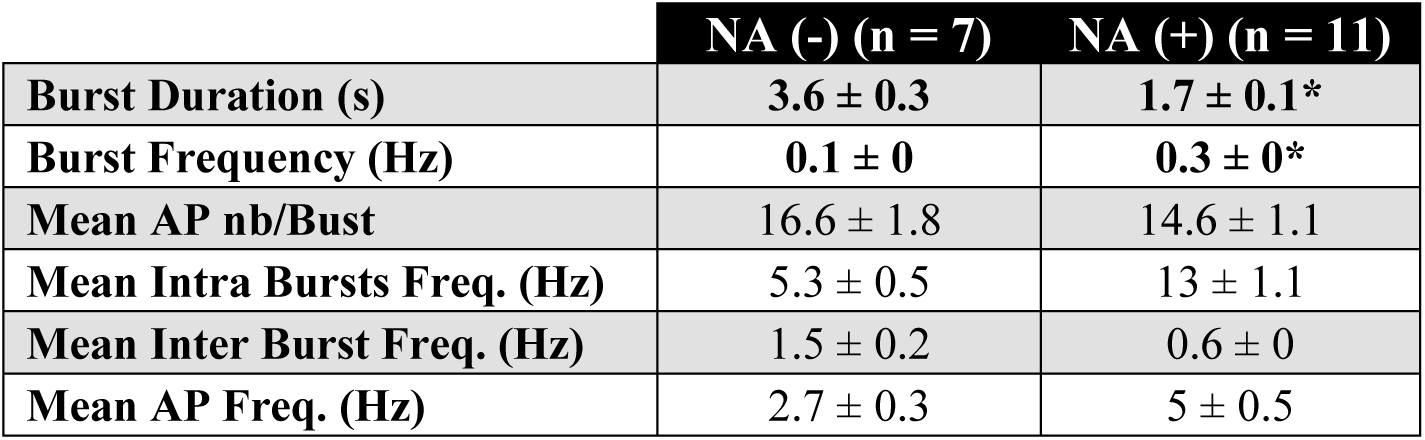
Bursting features of NA(-) and NA(+) neurons. Both NA(-) and NA(+) VLPO neurons exhibit burst firing activity with broadly similar properties, except for their burst duration and burst frequency that differ significantly depending on their response to NA application. *P < 0.05, Mann-Whitney test.

|  | NA (-) (n = 7) | NA (+) (n = 11) |
| --- | --- | --- |
| **Burst Duration (s)** | **$3.6 \pm 0.3$** | **$1.7 \pm 0.1^*$** |
| **Burst Frequency (Hz)** | **$0.1 \pm 0$** | **$0.3 \pm 0^*$** |
| **Mean AP nb/Burst** | $16.6 \pm 1.8$ | $14.6 \pm 1.1$ |
| **Mean Intra Bursts Freq. (Hz)** | $5.3 \pm 0.5$ | $13 \pm 1.1$ |
| **Mean Inter Burst Freq. (Hz)** | $1.5 \pm 0.2$ | $0.6 \pm 0$ |
| **Mean AP Freq. (Hz)** | $2.7 \pm 0.3$ | $5 \pm 0.5$ |

We next analyzed VLPO neurons recorded in the whole-cell configuration to explore how bursting relates to intrinsic membrane potential dynamics and neuronal type. Out of 24 neurons, 5 were inhibited in response to NA application, thus corresponding to putative sleep-promoting neurons. To further explore whether distinct subpopulations of bursting neurons were present in our sample, we examined intrinsic electrophysiological properties estimated immediately after achieving intracellular access as well as morphological characteristics estimated from infrared pictures of somata taken before recordings and 3-dimensional reconstructions of dendritic arbors obtained after intracellular biocytin filling and subsequent Neurolucida reconstructions (Materials and Methods).

A feature set of 34 parameters encompassing burst dynamics, as well as electrophysiological and morphological properties, was selected for unsupervised clustering (Materials and Methods). Ward’s method (Ward, 1963) was first applied to build a dendrogram representing the similarity between bursting neurons (Fig. 2A). Clusters generated by Ward’s method were corrected by applying the k-means algorithm to ensure that clusters occupied non-overlapping regions of the feature space (Dubourget et al., 2017; Perrenoud et al., 2012). Partitions in 2 to 6 clusters were considered (Fig. S1) (Tan et al., 2019). The total within-cluster sum of squared errors (SSE) and average silhouette value presented an elbow around 3 clusters (Fig. 2B), indicating that this number represents a good trade-off between model simplicity and explanatory power. When considering 3 groups, the k-means algorithm reassigned a total of 3 neurons from Ward’s dendrogram (Fig. 2C), resulting in a partition having a mean silhouette value of 0.205 (Fig. 2D).

**Figure 2.**
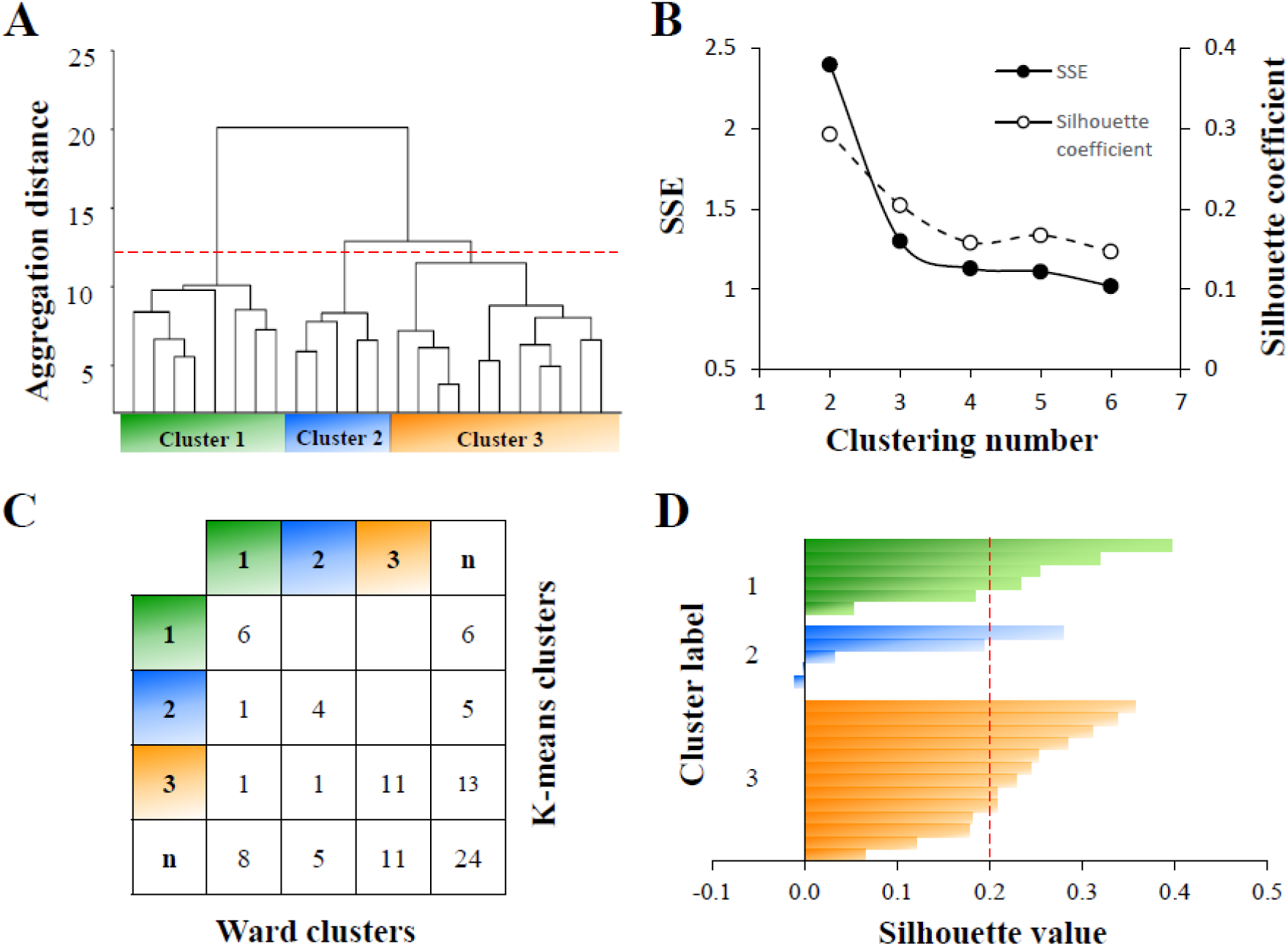
Unsupervised clustering of bursting VLPO neurons. **A**, Ward’s clustering of 24 bursting neurons. Individual cells are represented along the x-axis. The y-axis represents the average within-cluster linkage distance in a space of 34 electrophysiological and morphological characteristics. Three clusters: 1 (in green) and 2 (in blue), and 3 (in orange) were identified. **B**, SSE, and silhouette coefficients *versus* number of clusters. **C**, Clusters generated by Ward’s method in (A) were corrected using the clustering output generated by the *K-means* algorithm. **D**, The silhouette analysis was performed to assess the quality of the clustering (mean value of 0.203; red dashed line).

**Figure 3.**
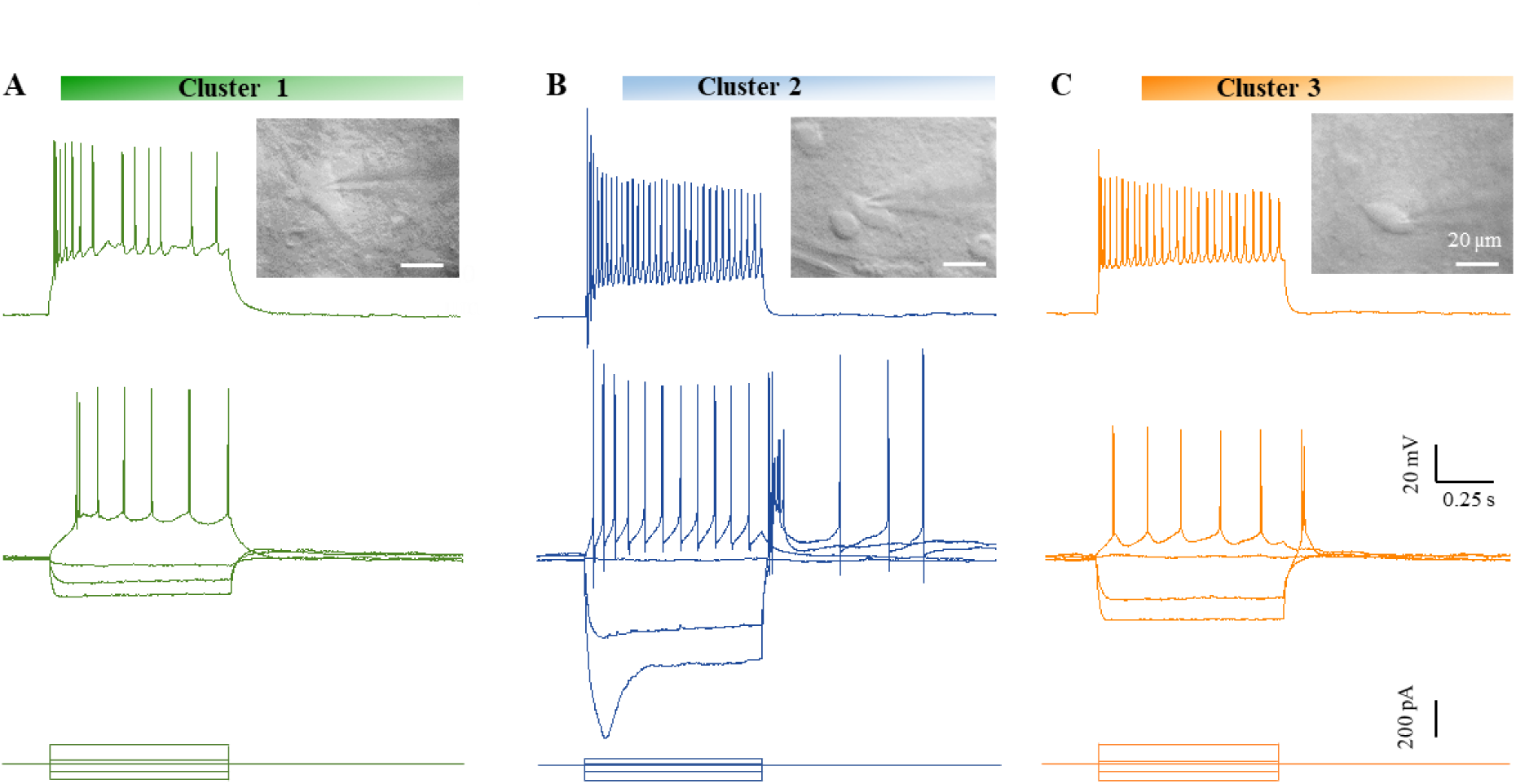
Passive membrane properties and Firing properties of the bursting VLPO neurons. A, Typical electrophysiological behavior of neurons from cluster 1 (green, *top*). Insert: Infrared image of the recorded neuron in whole-cell configuration. A strong depolarizing current (*top trace*) evoked a high and sustained firing. Responses to-100,-50, 0 pA and threshold current pulses (*middle and bottom traces*). **B** and **C**, same as in A but for a neuron from cluster 2 (blue) and cluster 3 (yellow).

Neurons in cluster 1 (n = 6/24) were all excited by NA (Fig. S2) and rarely displayed LTS or rebound from hyperpolarization (Table 2), consistent with a non-sleep-promoting neuron phenotype. These neurons had relatively low membrane resistance, high rheobases (Table 2) and frequently displayed a stuttering firing pattern (Fig. 2A, 3A). Bursting in these neurons was not accompanied by marked variations in the membrane potential called UP and DOWN states (Fig. 4A, Table 3). Bursts of action potentials were typically short and alternated rapidly with epochs of silence, recurring intrinsically with a periodicity of once every ∼200 ms (Fig. 4B, Table 3). Morphologically, cluster 1 neurons present large fusiform somata and well-developed dendritic arbor that extended significantly farther than in other clusters (Fig. 5, Table 4).

**Figure 4.**
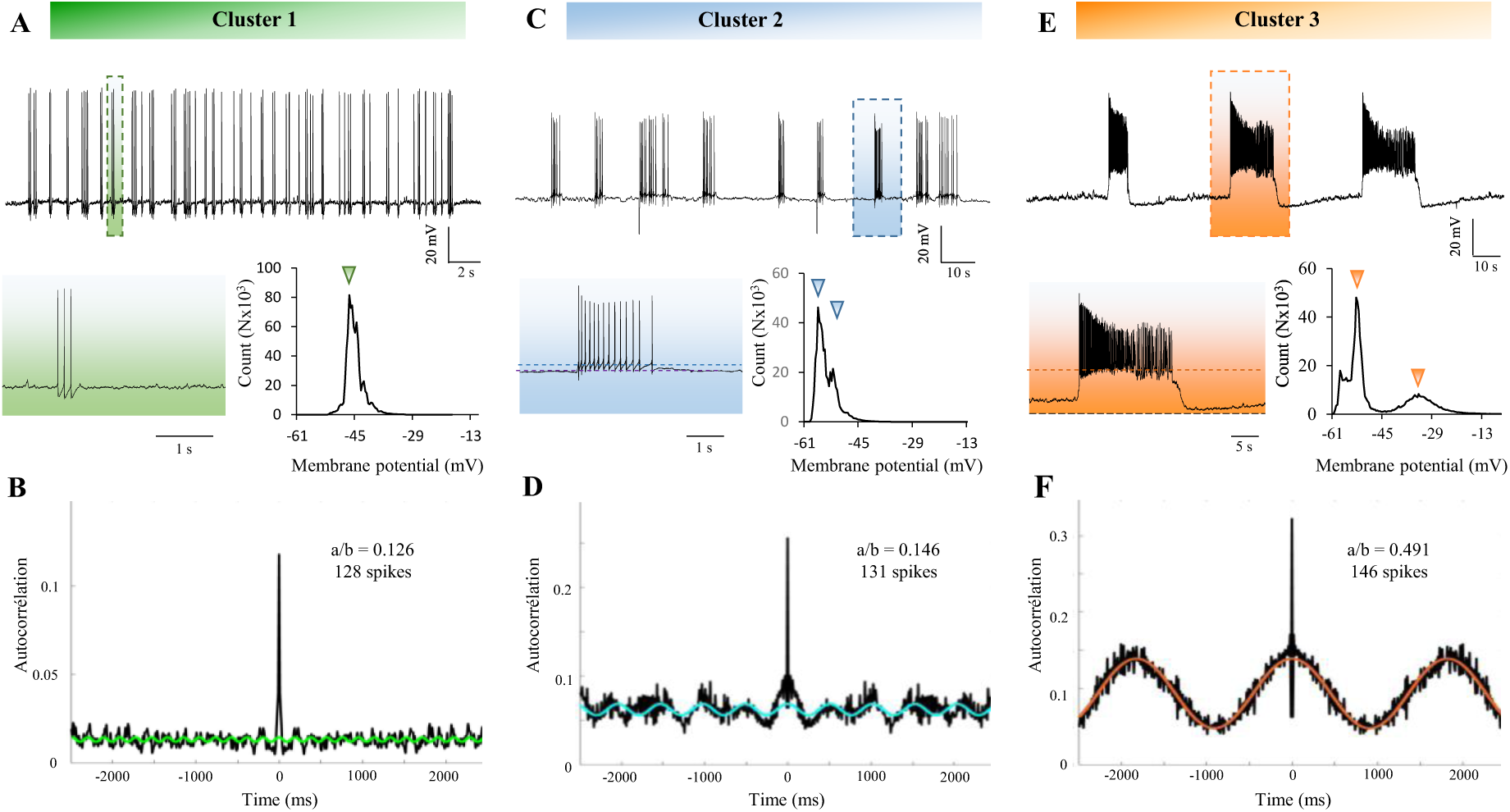
**Busting properties and passive membrane features**. **A**, Whole-cell recording of spontaneous bursting firing at the resting membrane potential (RMP) of-50 mV (*top*). Inset (green rectangle): magnified view of a single burst. Bottom right: histogram of membrane potential values for cluster 1 neurons, with the preferred membrane potential indicated by an arrow. **B**, Spiking auto-correlogram reveals a periodicity. **C-F**, same as in A-B but for a neuron from clusters 2 (in blue, with a RMP of-60 mV) and 3 (in orange, with a RMP of-50 mV).

**Figure 5.**
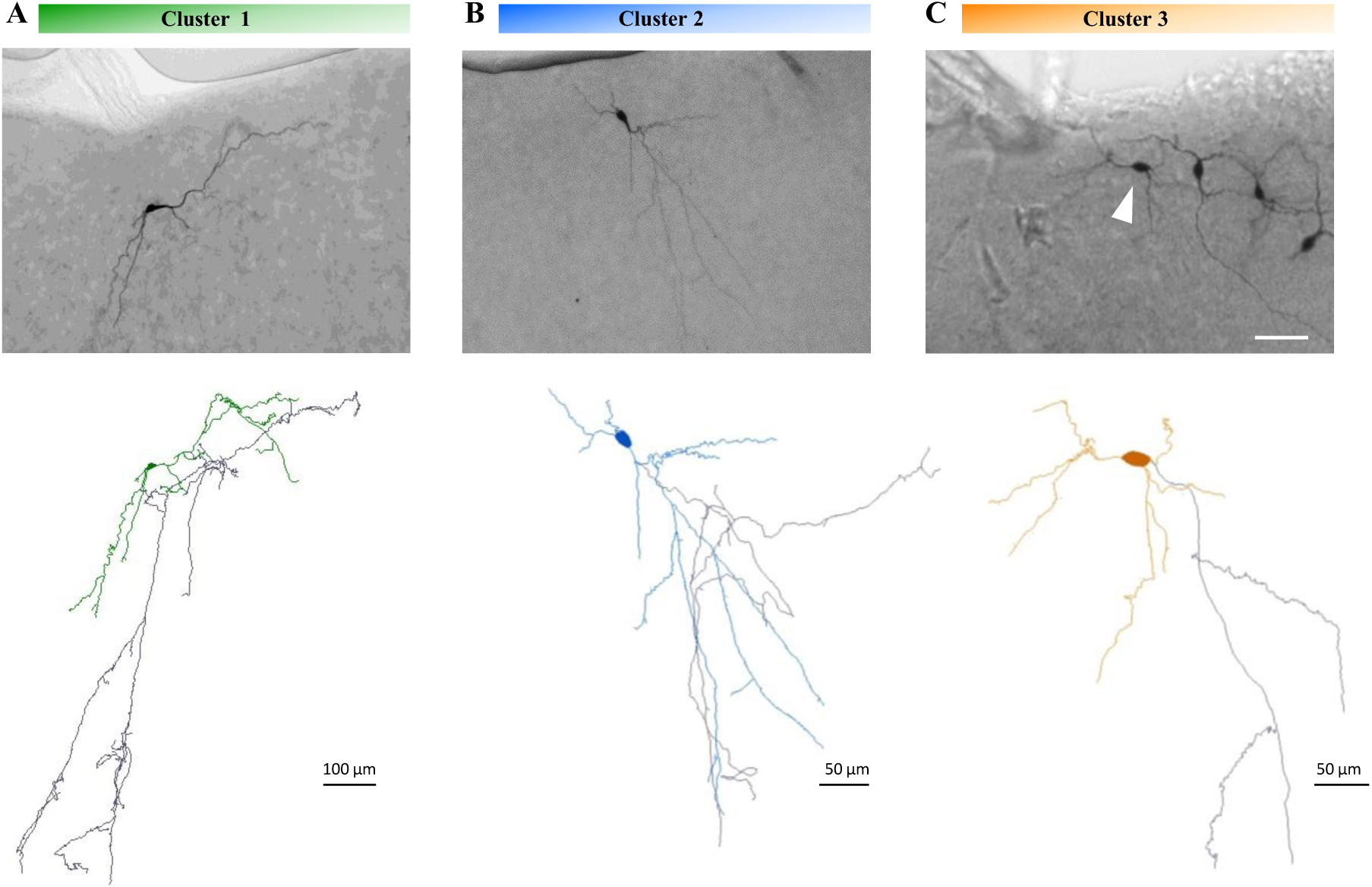
Neurolucida reconstructions of typical bursting neurons from the three clusters. **A**, Representative cluster 1 neuron: epifluorescence image of a biocytin-filled bursting VLPO neuron (*top*) and its Neurolucida reconstruction (*bottom*). Dendrites and soma are shown in green; the axon in grey. **B**, Same as in (A) for a cluster 2 neuron. **C**, Same as in (A) for a cluster 3 neuron. The white arrowhead indicates the recorded, biocytin-filled neuron.

**Table 2:**
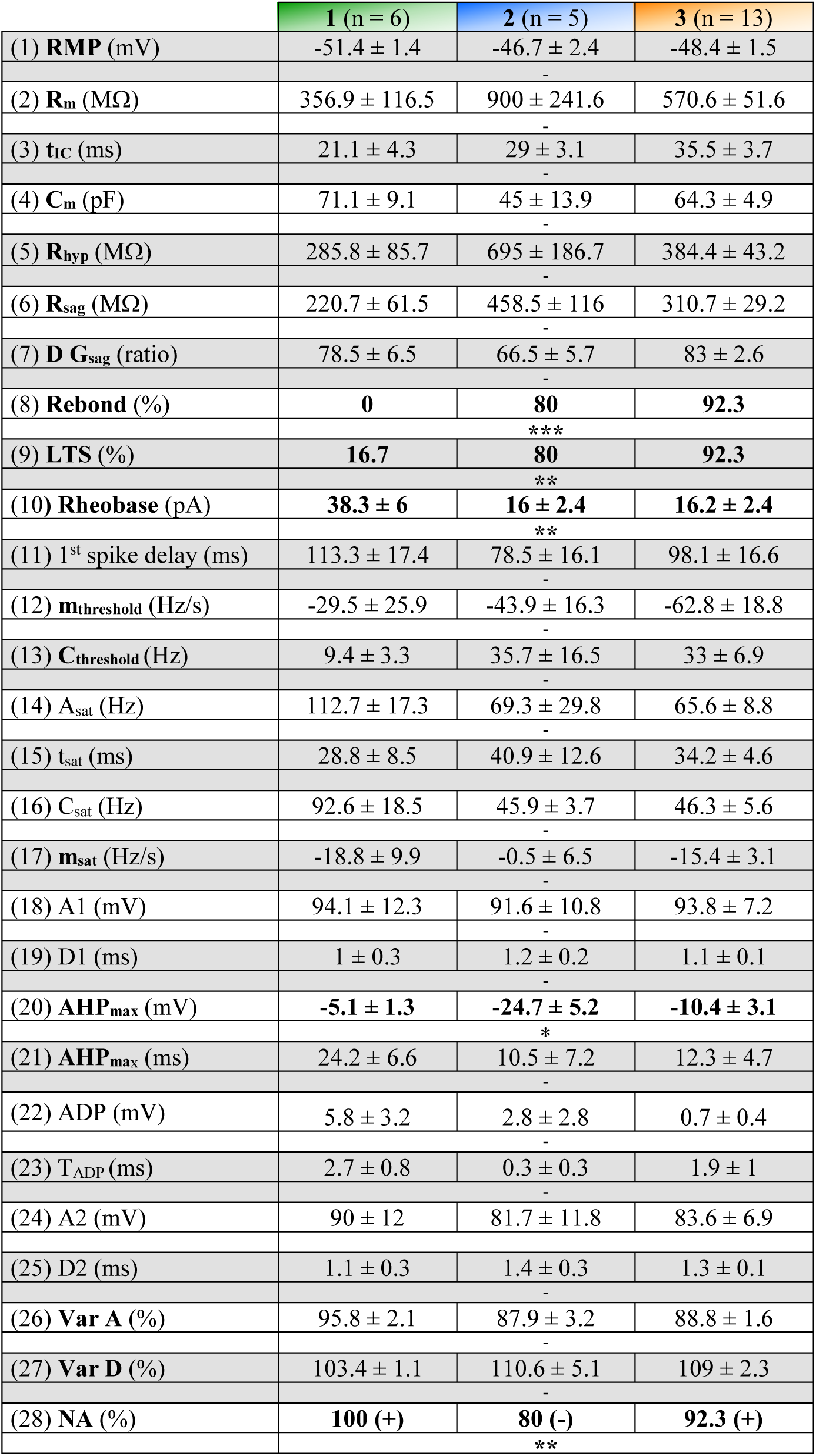
Electrophysiological properties of bursting VLPO neurons. Kruskal–Wallis test, followed by a Dunn’s post hoc test.

**Table 3:**
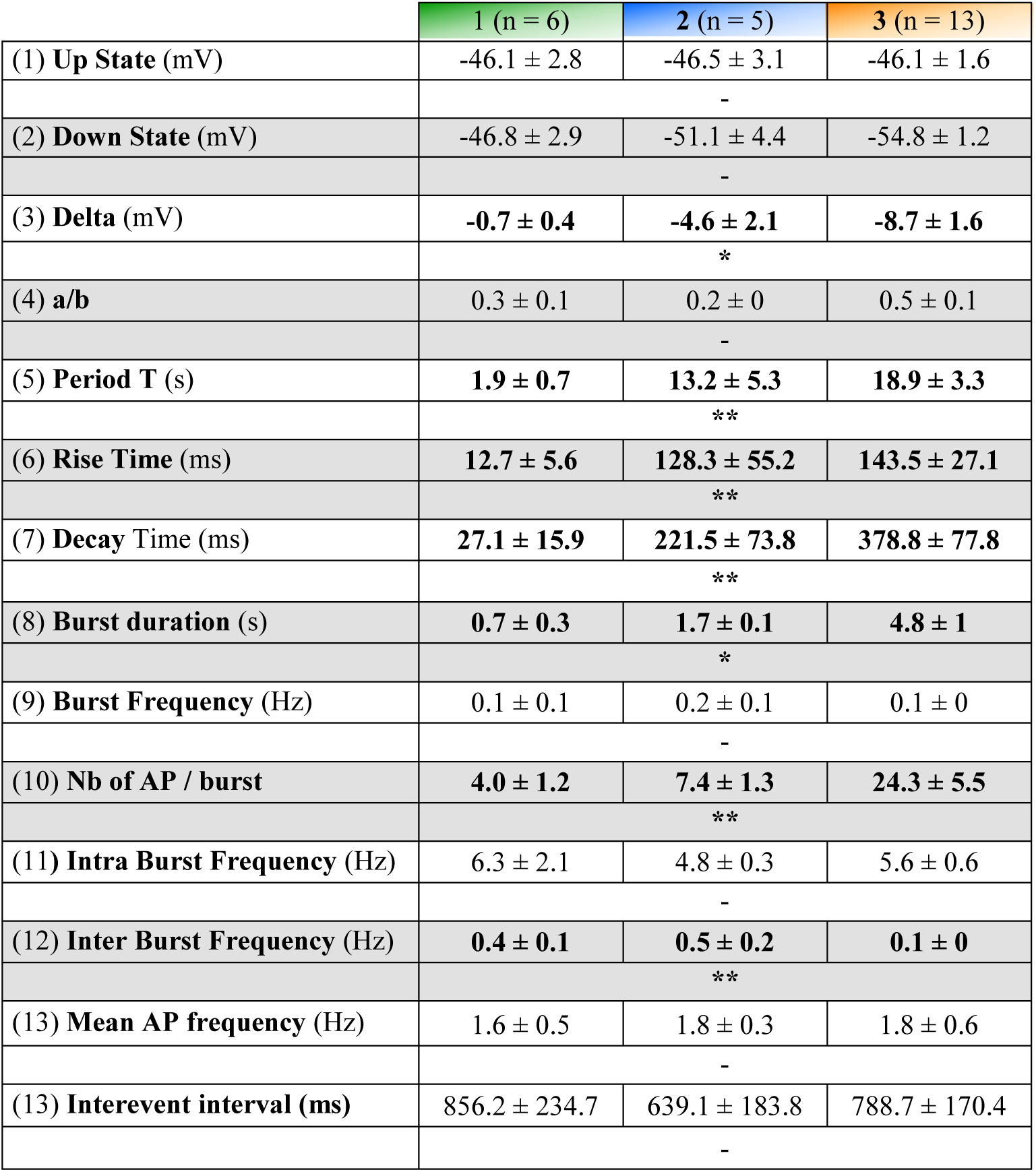
Bursting properties of VLPO neurons. Kruskal–Wallis test, followed by a Dunn’s post hoc test.

|  | 1 (n = 6) | 2 (n = 5) | 3 (n = 13) |
| --- | --- | --- | --- |
| (1) **Up State** (mV) | -46.1 ± 2.8 | -46.5 ± 3.1 | -46.1 ± 1.6 |
|  | - |  |  |
| (2) **Down State** (mV) | -46.8 ± 2.9 | -51.1 ± 4.4 | -54.8 ± 1.2 |
|  | - |  |  |
| (3) **Delta** (mV) | **-0.7 ± 0.4** | **-4.6 ± 2.1** | **-8.7 ± 1.6** |
|  | * |  |  |
| (4) **a/b** | 0.3 ± 0.1 | 0.2 ± 0 | 0.5 ± 0.1 |
|  | - |  |  |
| (5) **Period T** (s) | **1.9 ± 0.7** | **13.2 ± 5.3** | **18.9 ± 3.3** |
|  | ** |  |  |
| (6) **Rise Time** (ms) | **12.7 ± 5.6** | **128.3 ± 55.2** | **143.5 ± 27.1** |
|  | ** |  |  |
| (7) **Decay Time** (ms) | **27.1 ± 15.9** | **221.5 ± 73.8** | **378.8 ± 77.8** |
|  | ** |  |  |
| (8) **Burst duration** (s) | **0.7 ± 0.3** | **1.7 ± 0.1** | **4.8 ± 1** |
|  | * |  |  |
| (9) **Burst Frequency** (Hz) | 0.1 ± 0.1 | 0.2 ± 0.1 | 0.1 ± 0 |
|  | - |  |  |
| (10) **Nb of AP / burst** | **4.0 ± 1.2** | **7.4 ± 1.3** | **24.3 ± 5.5** |
|  | ** |  |  |
| (11) **Intra Burst Frequency** (Hz) | 6.3 ± 2.1 | 4.8 ± 0.3 | 5.6 ± 0.6 |
|  | - |  |  |
| (12) **Inter Burst Frequency** (Hz) | **0.4 ± 0.1** | **0.5 ± 0.2** | **0.1 ± 0** |
|  | ** |  |  |
| (13) **Mean AP frequency** (Hz) | 1.6 ± 0.5 | 1.8 ± 0.3 | 1.8 ± 0.6 |
|  | - |  |  |
| (13) **Interevent interval** (ms) | 856.2 ± 234.7 | 639.1 ± 183.8 | 788.7 ± 170.4 |
|  | - |  |  |

**Table 4.** Somatic properties of bursting neurons on infrared images before patch-clamp recordings. n, number of cells; Kruskal–Wallis test.

|  | 1 (n = 6) | 2 (n = 5) | 3 (n = 13) |
| --- | --- | --- | --- |
| (1) **Cell body area** ($\mu\text{m}^2$) | 253.1 ± 52.8 | 181.5 ± 32.5 | 193.9 ± 13.2 |
| (2) **Perimeter** ($\mu\text{m}$) | 68.6 ± 9.3 | 53.8 ± 5.5 | 56.9 ± 3.3 |
| (3) **Form factor** | 0.7 ± 0.1 | 0.8 ± 0 | 0.8 ± 0 |
| (4) **Feret max** ($\mu\text{m}$) | 26.1 ± 5.2 | 21 ± 2.6 | 21.8 ± 1.4 |
| (5) **Feret min** ($\mu\text{m}$) | 13.1 ± 1.5 | 11.8 ± 1.3 | 13.2 ± 0.6 |
| (6) **Aspect ratio** | 2 ± 0.4 | 1.9 ± 0.3 | 1.6 ± 0.1 |
| (7) **Solidity** | 1 ± 0 | 1 ± 0 | 0.9 ± 0 |
| (8) **Convexity** | 0.9 ± 0 | 1 ± 0 | 1 ± 0 |
| (9) **Roundness** | 1.5 ± 0.2 | 1.3 ± 0.1 | 1.3 ± 0.1 |
| (10) **Compactness** | 1.2 ± 0.1 | 1.1 ± 0 | 1.2 ± 0 |

**Table 5:** Neurolucida-based dendritic morphology analysis. n, number of cells; Kruskal– Wallis test, followed by a Dunn’s post hoc test. Asterisks indicate values significantly different in one cluster vs. all others.

|  | 1 (n = 4) | 2 (n = 4) | 3 (n = 7) |
| --- | --- | --- | --- |
| (1) Nb of dendrites | 4.5 ± 0.2 | 4 ± 0.4 | 3.9 ± 0.3 |
|  | - | - | - |
| (2) Dendritic length (µm) | 3733.3 ± 405.8 | 2185.6 ± 654.5 | 2184.9 ± 266.4 |
|  | - | - | - |
| (3) Dendritic surface (µm<sup>2</sup>) | 10118.5 ± 1734.8 | 5706.1 ± 1507.5 | 5410.8 ± 693.4 |
|  | - | - | - |
| (4) Dendritic volume (µm<sup>3</sup>) | 3112.5 ± 697.4 | 1685 ± 413 | 1494.4 ± 236.3 |
|  | - | - | - |
| (5) Length/surface | 0.4 ± 0 | 0.4 ± 0 | 0.4 ± 0 |
|  | - | - | - |
| (6) Highest order segment | 7.3 ± 0.5 | 6.5 ± 1.4 | 6.1 ± 0.5 |
|  | - | - | - |
| (7) Dendritic tortuosity | 1.4 ± 0 | 1.4 ± 0 | 1.3 ± 0 |
|  | - | - | - |
| (8) Nodes | 24.8 ± 1.9 | 18 ± 7.6 | 13.7 ± 1.7 |
|  | - | - | - |
| (9) Dendritic planar angle | 42.9 ± 2 | 44.6 ± 3.2 | 42.7 ± 2.5 |
|  | - | - | - |
| (10) Dendritic spline angle | 47.9 ± 1.9 | 48.6 ± 4.4 | 47.1 ± 2.5 |
|  | - | - | - |
| (11) Dendritic sholl (100) | 1175.3 ± 224.3 | 913.3 ± 247.6 | 803.3 ± 77.8 |
|  | - | - | - |
| (12) Dendritic sholl (200) | 1166.6 ± 196.9 | 809.7 ± 358.5 | 709.5 ± 134.7 |
|  | - | - | - |
| (13) Dendritic sholl (300) | **874.8 ± 95.1** | **254.3 ± 124.7** | **343.8 ± 81.2** |
|  | - | * | - |
| (15) Kdim | 1.1 ± 0 | 1.1 ± 0 | 1.1 ± 0 |
|  | - | - | - |
| (16) Spine 1 | 13.5 ± 9.7 | 6 ± 2.9 | 10 ± 5.2 |
|  | - | - | - |
| (17) Spine 2 | 18.8 ± 8.4 | 33.8 ± 12.5 | 39.9 ± 10.5 |
|  | - | - | - |
| (18) Spine 3 | 38.3 ± 25.8 | 49.3 ± 22.8 | 47.7 ± 8.9 |
|  | - | - | - |
| (19) Spine 4 | 30.5 ± 24.1 | 66.5 ± 35.3 | 33.4 ± 4.5 |
|  | - | - | - |
| (20) Spine 5 | 16.3 ± 5.3 | 15.5 ± 10.8 | 15.3 ± 5.4 |
|  | - | - | - |
| (21) Spine 6 | 29.3 ± 13.1 | 7 ± 6 | 22 ± 9.7 |
|  | - | - | - |
| (22) Spine 7 | 3 ± 1.2 | 11 ± 9.8 | 7.6 ± 5.6 |
|  | - | - | - |
| (23) Spine 8 | 11.3 ± 8.6 | 1.8 ± 1.6 | 0.9 ± 0.6 |
|  | - | - | - |
| (24) Spine density | 0 ± 0 | 0.1 ± 0 | 0.1 ± 0 |
|  | - | - | - |

Cluster 2 neurons (n = 5/24) were mostly inhibited by NA (Fig. S2) and displayed prominent LTS and rebound following hyperpolarization (Table 2), consistent with a sleep-promoting phenotype. Accordingly, these neurons presented a high membrane resistance, a pronounced delayed rectification in response to hyperpolarizing current pulses, and marked AHPs (Fig. 3B, Table 2). Up and down states in cluster 2 neurons occurred at intermediate frequencies, recurring intrinsically with a periodicity of once every ∼500 ms and were accompanied by a ∼5 mV switching between the bimodal distribution of the membrane potential (Fig. 4 C-D, Table 3). Morphologically, these neurons had relatively small fusiform somata and medium-sized dendritic arbors (Fig. 5B, Table 4).

Cluster 3 neurons were the most prevalent bursting subtype (n = 13/24). They exhibited LTS and rebound from hyperpolarization (Fig. 3C, Table 2). However, they were excited by NA, indicating a non-sleep-promoting phenotype (Table 2). Their intrinsic electrophysiological properties represented an intermediate between the properties of clusters 1 and 2. Yet, their bursting properties were distinctive: bursts of action potential were long, occurred at low frequencies, recurring with a periodicity of once every ∼2 s, and were accompanied by marked transitions between UP and DOWN state of the membrane potential of ∼9 mV (Fig. 4E-F, Table 3) (Bal and McCormick, 1993; Blethyn et al., 2006; Brunton and Charpak, 1997; Domich et al., 1986; M. V Sanchez-Vives and McCormick, 2000; Steriade et al., 1993) Morphologically, cluster 3 neurons had medium-sized, ovoid cell bodies and dendritic arbors typically extending parallel to the pia (Fig. 5C, Table 4-5).

To gain insight into the mechanism underlying bursting, we examined responses to depolarizing current steps (10, 20, 30 and 40 pA; Fig. 6). In clusters 2 (n = 3) and 3 (n = 7), increasing current injections tended to increase the number of AP per burst and the burst frequency, with the largest steps often switching neurons in a state of tonic firing (Fig. 6B C). Interestingly, however, cluster 1 neurons tended to maintain a stuttering pattern upon current injection, and the strongest current step tended to reduce overall firing, possibly reflecting saturation (n = 5, Fig. 6A).

**Figure 6.**
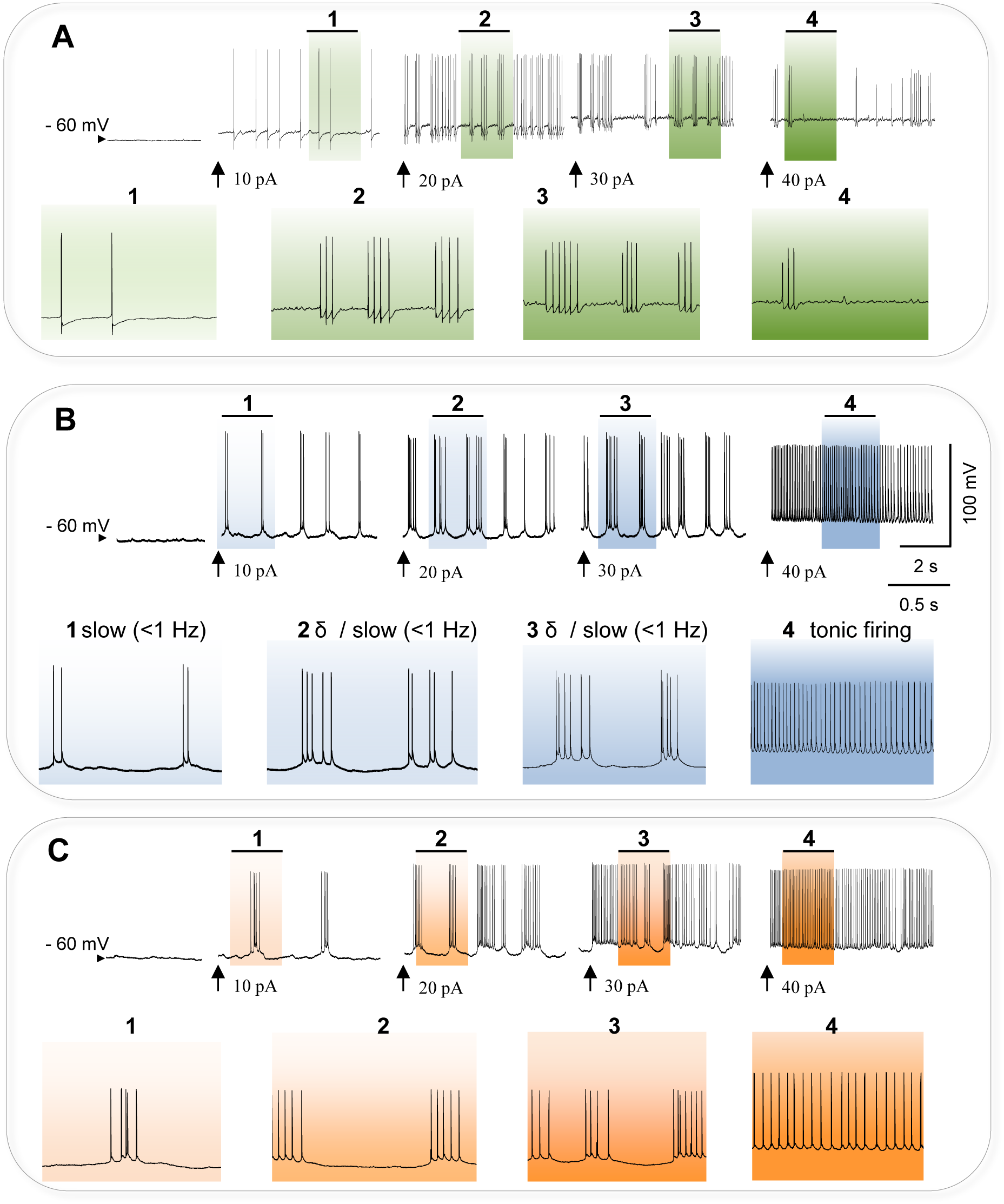
The slow (<1 Hz) oscillation forms part of a continuum of activity in VLPO neurons. **A**, Whole-cell recording in current-clamp mode of the oscillatory activity. A typical VLPO neuron from cluster 1 is held at different membrane potential as indicated by arrows. Injections of current pulses progressively led to continuous tonic firing. Sections marked above are expanded below. **B, C,** Same as in A, but for neurons from clusters 2 and 3, respectively.

We next tested TTX on spontaneous bursting activity. As expected, TTX abolished spiking in all neurons. Interestingly, however, oscillations of the membrane potential persisted in all neurons except for 2 neurons from cluster 3 (Fig. 7 A-D). This strongly suggests that oscillations of the membrane potential are generated intrinsically in those neurons. In the remaining 2 neurons from cluster 3, TTX completely abolished the oscillatory UP and DOWN dynamics of the membrane potential, suggesting that bursting was induced by synaptic currents.

**Figure 7.**
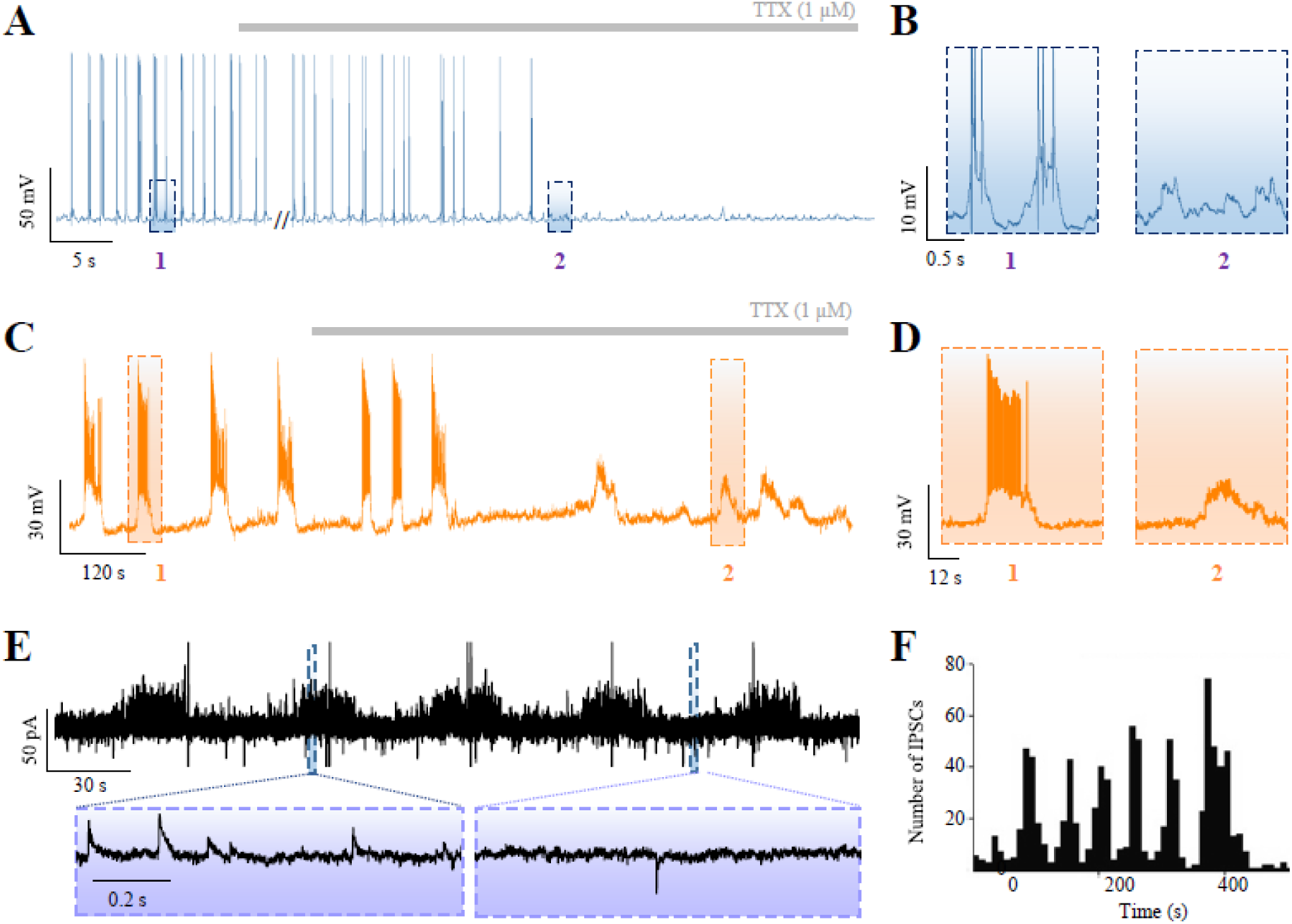
TTX-resistant membrane oscillations and burst-like sIPSCs in VLPO neurons. A-D, Under TTX, neurons from clusters 2 (n = 2) and 3 (n = 5) continue to exhibit spontaneous membrane-potential oscillations. Inset: High magnification of the UP state indicated by a red dotted rectangle. **E**, In 4 out of 66 neurons, recordings of spontaneous inhibitory postsynaptic currents (sIPSCs) revealed burst-like episodes. Left: intraburst epoch (red dotted rectangle; high sIPSC frequency). Right: interburst epoch (red dotted rectangle; low sIPSC frequency). **F** Histogram of sIPSC event counts for the recording shown in (E).

To evaluate a synaptic contribution more directly, we examined voltage-clamp recordings of spontaneous excitatory and inhibitory postsynaptic currents (sEPSCs and sIPSCs) obtained in 66 neurons out of 289 whole-cell recordings (Dubourget et al., 2017). We identified 4 neurons that presented synchronized bursts of IPSCs (Fig. 7E, F). Intraburst sIPSCs frequency (2.75 ± 0.36 Hz) was significantly higher than the interburst frequency of (0.30 ± 0.09 Hz; *n* = 4; *P* <0.029; Mann-Whitney *U*-test). All these neurons were NA(-), and 3 out of 4 were endowed with an LTS. The combined observation of UP and DOWN-state and bursts of sIPSCs in neighboring neurons suggests that intrinsically bursting neurons can propagate slow rhythmic activity to NA(-) sleep-promoting neurons of the VLPO through inhibitory synaptic inputs.

## Discussion

Here, we report that VLPO neurons can exhibit intrinsically generated rhythmic bursting activity. Spontaneous bursting was screened in a database of 393 *ex-vivo* whole-cell and loose cell-attached recordings and occurred in an estimated 12.47% of VLPO neurons. Analyses of loose-patch recordings revealed bursting in both NA(-) putative sleep-promoting neurons and NA(+) putative wake-active neurons. Unsupervised clustering performed on 24 whole-cell recordings confirmed that bursting behavior in the VLPO occurs in NA(-) putative sleep-promoting neurons and discriminated two NA(+) putative wake-active neuron subtypes, respectively displaying fast and slow bursting dynamics. Interestingly, oscillations of the membrane potential persist in the presence of TTX in most tested neurons, indicating that the underlying rhythm is predominantly intrinsic rather than synaptically driven. Collectively, these findings suggest that the VLPO contributes to both the generation and the propagation of slow rhythmic activity.

To our knowledge, our study constitutes the first report of rhythmic bursting activity in VLPO neurons. The lack of a previous report likely reflects the low probability of observing spontaneous bursting in ex vivo slices, and the need for large datasets. Here, the study of spontaneous bursting was conducted in parallel with other studies of the properties of VLPO neurons (Dubourget et al., 2017; Sangare et al., 2016; Scharbarg et al., 2016; C. Varin et al., 2015), making it possible to sample a large dataset.

### Intrinsic oscillation dynamics and reinforcing GABAergic pacing

On one hand, we observed that in most recorded neurons, slow UP/DOWN oscillations of the membrane potential persisted in TTX, indicating an intrinsic generator. On the other hand, we observed rhythmically modulated GABAergic inputs in neighboring VLPO neurons. These inhibitory events occurred at the **s**ame rhythm as the intrinsic oscillation and could therefore entrain and reinforce the rhythmic discharge. A parsimonious interpretation is that intrinsically oscillating VLPO neurons rhythmically inhibit nearby sleep-promoting neurons, thereby sharpening or amplifying their slow bursting through phase-locked IPSCs, consistent with synchronous sIPSC bursts observed in NA(–) cells. These inhibitory inputs could arise from neighboring bursting VLPO neurons, effectively coupling intrinsic and synaptic drivers of the slow rhythm.

### Cluster-specific mechanisms

Understanding the function of rhythmic bursting in the VLPO might require identifying and modulating the ion conductance involved in its initiation and maintenance. It is worth noting that rhythmic bursting presented distinct temporal characteristics in neurons from clusters 1 and 3, suggesting that it might rely on distinct mechanisms in these two groups. Accordingly, while current injection tended to abolish bursting and induced a state of continuous firing in neurons from clusters 2 and 3, bursting was maintained in neurons from cluster 1. Cluster 1 Na(+) and stuttering neurons tend to maintain a relatively depolarized state, and their bursting appeared to result from failures of spike initiation (Fig. 3A, 6A). This pattern is reminiscent of a ‘stuttering’ firing, which has been described in cortical interneurons and depends on D-type K^+^ currents (Golomb et al., 2007; Markram et al., 2004; Tepper and Bolam, 2004). By contrast, bursting in neurons from clusters 2 and 3 seems to rely on intrinsically generated oscillations of the membrane potential (Fig. 3B-C, 6B-C). Similar dynamics have been described in cortical layer 5 (Beltramo et al., 2013; Sanchez-Vives and McCormick, 2000; Steriade et al., 1993) and 6b (WengerCombremont et al., 2016), shown to depend on persistent non-inactivating voltage sodium currents (Le Bon-Jego and Yuste, 2007). However, such conductance is TTX-sensitive (Storm, 1988). Thus, VLPO membrane potential oscillations likely depend on TTX-insensitive conductance acting similarly. Interestingly, neurons from clusters 2 and 3 displayed LTS, which is thought to depend on T-type calcium currents (Crunelli et al., 2005, 1989). These currents are capable of inducing spontaneous depolarization of the membrane potential (Sherman, 2001).

While T-type or similarly acting currents may induce depolarization, oscillations also require counteracting hyperpolarizing mechanisms. As depolarization does not rely on TTX-sensitive voltage-gated Na^+^ channels, hyperpolarization is, in turn, unlikely to require Na^+^-activated K^+^ currents (Bhattacharjee and Kaczmarek, 2005; Kim and McCormick, 1998). In contrast, Ca^2+^ activated K^+^ currents are plausible contributors (Vergara et al., 1998).

### From single-cell pacemakers to local microcircuits

Electrical synapses being particularly effective for generating synchrony within networks of inhibitory neurons (Landisman et al., 2002; Long et al., 2004), gap-junctions could shape firing patterns within the VLPO, as suggested in the thalamus (Blethyn et al., 2006). In addition, dynamic interactions between neurons and glial cells may modulate information processing within the VLPO, as it was reported in the olfactory bulb (Roux et al., 2015). Together with the phase-locked GABAergic inputs seen in NA(–) neurons, these mechanisms could be hybrid: intrinsic oscillators providing the UP state drive, while local inhibitory coupling (and possibly gap junctions or astroglial mechanisms) phase-aligning and amplifying the rhythm within VLPO.

### Generalization across the hypothalamus

Interestingly, our results are reminiscent of data obtained in other regions of the hypothalamus. Robust evoked bursts of mIPSCs were indeed also recorded in hypothalamic magnocellular neurons (Popescu et al., 2010). In contrast, synchronized EPSPs have been described in oxytocin neurons in organotypic hypothalamic cultures (Israel et al., 2003). These pieces of evidence suggest that the ability to produce rhythmic bursting is widespread in the hypothalamus and that the hypothalamic network might shape the synchronization of spontaneous bursting firing in putative sleep-promoting neurons through bursting afferent volleys of IPSCs. Overall, bursting activity in the VLPO is likely governed by multiple interacting mechanisms, as in other brain regions (Beltramo et al., 2013; Le Bon-Jego and Yuste, 2007; Sanchez-Vives and McCormick, 2000; Steriade et al., 1993). Future work will thus have to disentangle these mechanisms to clarify the role of bursting activity in VLPO neurons.

### *In vivo* relevance

Although rhythmic activity has not been explicitly reported in *in vivo* recordings of sleep-active VLPO neurons, examination of their discharge reveals that such neurons often fire in bursts (Alam et al., 2014, 2004, 1999; Szymusiak et al., 1998; Takahashi et al., 2009). These bursting patterns may therefore reflect an underlying rhythmic process contributing to sleep induction and maintenance. However, it remains unknown whether VLPO neurons burst synchronously or maintain coordinated rhythmicity over time. In fact, it has already been shown in the hypothalamus that burst frequency varies considerably across neurons and even within individual cells during prolonged recordings (Takahashi et al., 2009), suggesting a flexible modulation rather than a fixed pacemaker rhythm.

It is well established that the firing of sleep-active VLPO neurons is modulated by sleep pressure (Alam et al., 2014), intrinsic factors such as adenosine (Alam et al., 1999; Gallopin et al., 2005; Gvilia et al., 2006), and synaptic inputs from interconnected regions (Chou et al., 2002; Chung et al., 2017; Walter et al., 2019). Within this modulatory framework, intrinsic oscillations may shape VLPO burst firing and its dynamic entrainment during sleep. These oscillations could act as a local coordinating signal, rhythmically inhibiting neighboring NA(–) neurons and promoting clustered GABA/galanin output from the VLPO, imposing a rhythmic inhibition on downstream wake-promoting centers such as the LC, TMN, and LH. The infraslow NA/LC fluctuations reported during NREM sleep by Lüthi and colleagues may reflect a periodic disinhibition of the LC driven by rhythmic inhibition within the VLPO (Osorio-Forero et al., 2025). Future *in vivo* recordings and causal manipulations of VLPO rhythmicity will be critical to determine whether these intrinsic oscillations represent a local microcircuit mechanism or a distributed pacemaker signal in the regulation of sleep dynamics.

## Acknowledgments

This work was supported by the Centre National de la Recherche Scientifique (CNRS), the French Institute of Health and Medical Research (Inserm). We thank all members of the animal house facility at ESPCI.

---

**Supplementary Figure 1.**
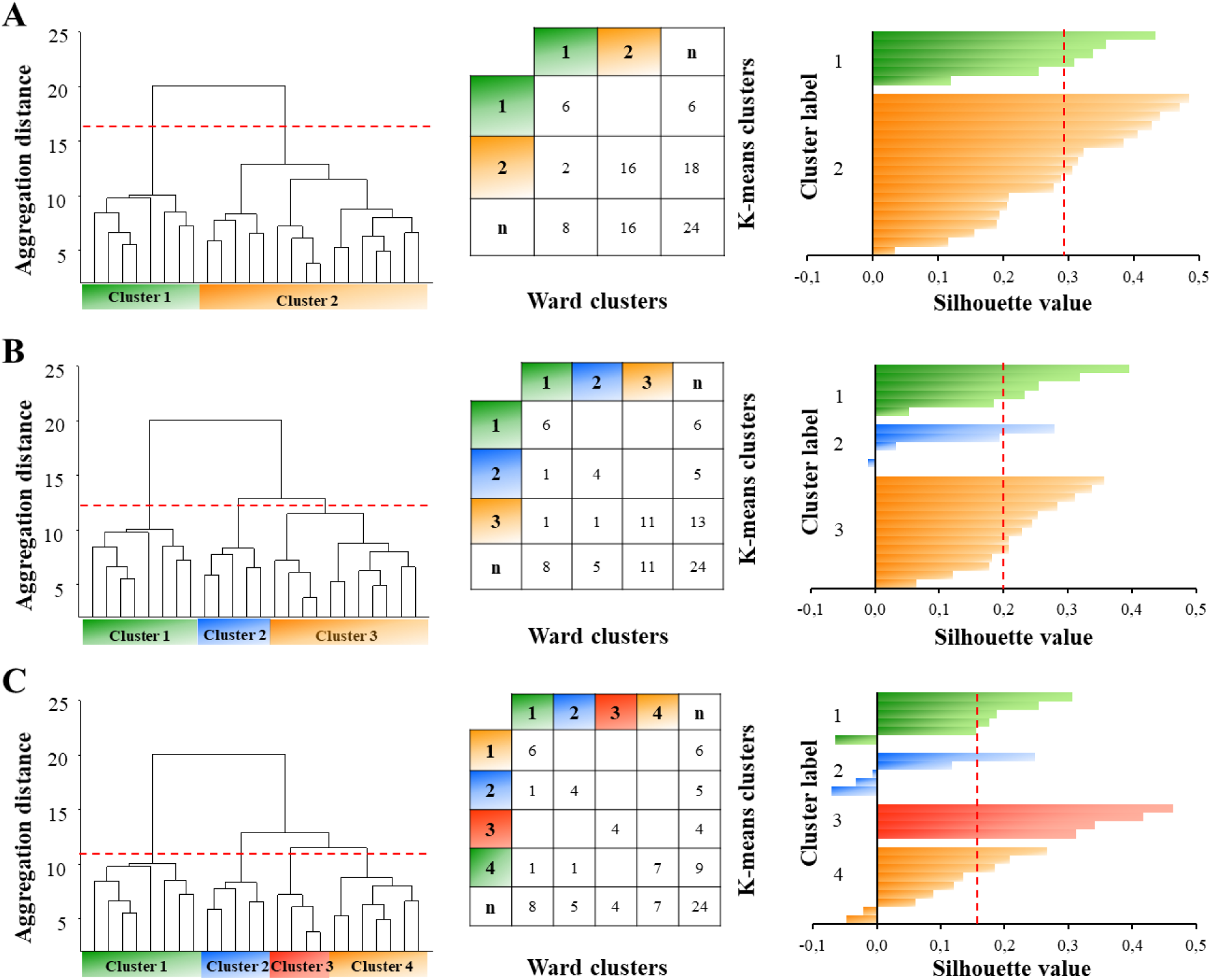
Unsupervised clustering of 24 bursting VLPO neurons. **A**, Ward’s clustering of 24 bursting neurons. Individual cells are represented along the x-axis. Same as in Fig. 2, with only two clusters: 1 (in green) and 2 (in orange) (*left*). Clusters generated by Ward’s method in (A) were corrected using the clustering output generated by the *K-means* algorithm (*middle*). The silhouette analysis was performed to assess the quality of the clustering (mean value indicated by the red dashed line). **B** and **C**, same as in A, with 3 and 4 clusters respectively.

**Supplementary Figure 2.**
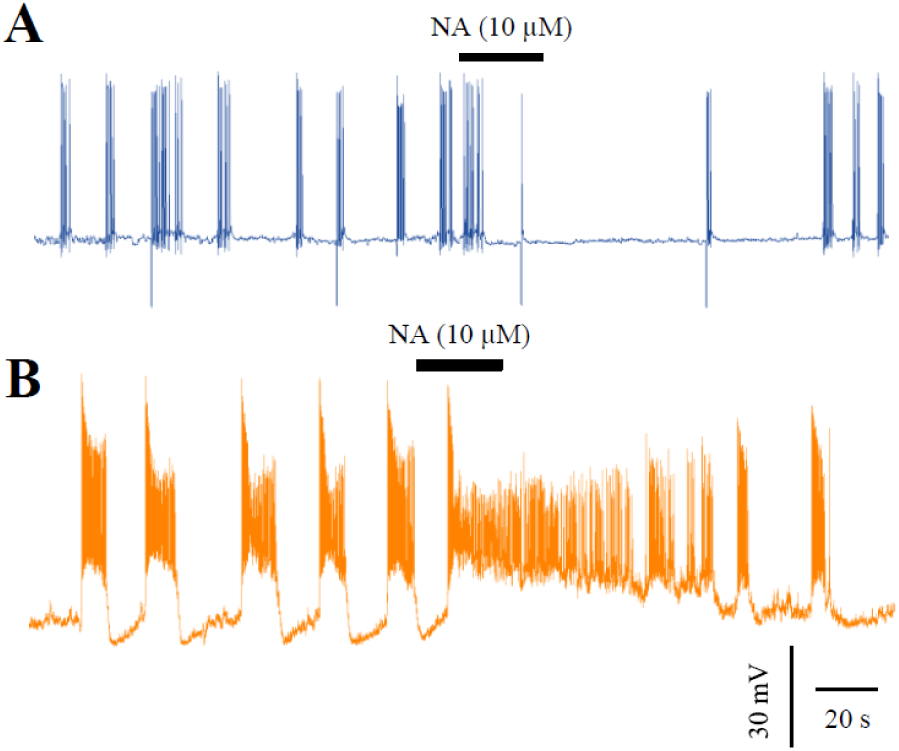
Pharmacological identification of sleep-promoting neurons. A, inhibitory effect of bath-applied (30 s) noradrenaline (NA, 10 µM). **B**, excitatory effect of bath-applied (30 s) noradrenaline (NA, 10 µM).

